# Whole protein sequencing and quantification without proteolysis, terminal residue cleavage, or purification: A computational model

**DOI:** 10.1101/2024.03.13.584825

**Authors:** G. Sampath

## Abstract

Sequencing and quantification of whole proteins in a sample without separation, terminal residue cleavage, or proteolysis are modeled computationally. Similar to recent work on DNA sequencing (*PNAS* **113**, 5233–5238, 2016), a high-volume conjugate is attached to every instance of amino acid (AA) type AA_i_, 1 ≤ i ≤ 20, in an unfolded whole protein, which is then translocated through a nanopore. From the volume excluded by 2L residues in a pore of length L nm (a proxy for the blockade current), a partial sequence containing AA_i_ is obtained. Translocation is assumed to be unidirectional, with residues exiting the pore at a roughly constant rate of ∼1/μs (*Nature Biotechnology* **41**, 1130–1139, 2023). The blockade signal is sampled at intervals of 1 μs and digitized with a step precision of 70 nm^3^; the positions of the AA_i_s are obtained from the positions of well-defined quantum jumps in the signal. This procedure is applied to all 20 standard AA types, the resulting 20 partial sequences are merged to obtain the whole protein sequence. The complexity of subsequence computation is O(N) for a protein with N residues. The method is illustrated with a sample protein from the human proteome (Uniprot id UP000005640_9606). A mixture of M’ protein molecules (including multiple copies) can be sequenced by constructing an M’ × 20 array of partial sequences from which proteins occurring multiple times are first isolated and their sequences obtained separately. The remaining M singly-occurring molecules are detected from M disjoint paths through the 20 columns of the reduced M × 20 array. Detection complexity is O(M^20^), which is nominally in polynomial time but practical only for small M; to use this method a sample may be subdivided into subsamples down to this level. Quantification of proteins can be done by sorting their computed sequences on the sequence strings and counting the number of duplicates. The possibility of translating this procedure into practice and related implementation issues are discussed.

## 1. Protein sequencing in the bulk and at the single molecule level

The primary sequence of a protein, the sequence of amino acids (AAs) that make up its primary structure, is central to practice in laboratory procedures in analytical chemistry and medical diagnostics and to studies in structural biology and biological modeling. The earliest known method of protein sequencing is Edman degradation [1], which is limited in its scope but continues to be used in the laboratory on a small scale. A second well-known method that is somewhat more prevalent is gel electrophoresis [2]. The most widely used method is mass spectrometry (MS) [3,4], a *de facto* industry standard that is undergoing continual improvement. All three methods are premised on the availability of bulk-sized samples. Unlike with DNA sequencing, which has transitioned from Sanger sequencing in the bulk to high-throughput Next Generation Sequencing methods that can work with sparse samples, in part because of the ability of PCR-based methods to amplify low-volume samples, there is no efficient way to amplify a protein sample. Protein sequencing is further hampered by the varying levels of electrical charge carried by individual residues, which can be positively or negatively charged or neutral, which makes their transport properties hard to predict or model. Discrimination among them is more difficult, in part because there are 20 individual AA types to identify (compared with 4 bases in DNA), many of which are very similar in their physical and chemical properties.

In its current state of development MS requires tens of thousands to a million or more copies of a protein. In many situations, such as single cell level analysis, this level of copy number is not available [5,6]. Ideally a protein sequencing method must be able to sequence a protein from a single copy or perhaps a few tens of copies. Such single molecule methods of protein sequencing (SMPS), have been pursued over the last decade or two, they continue to be a focus of research laboratories [7]. One such method is based on electrophoretically translocating a single protein molecule through a nano-sized pore in an electrolytic cell (e-cell) under the influence of an electric field [8]. During its passage through the pore the normal electrolytic current that flows in the absence of an analyte (which may be a polymer like DNA, RNA, or protein) experiences a drop or blockade, the size of which may be related to the monomer in some well-defined manner. Unlike nanopore DNA sequencing nanopore protein sequencing is yet to make significant strides toward reliable sequencing for research or for use in the field.

For recent reviews of protein sequencing, including SMPS, see [9,10]. Methods on the horizon include ongoing NIH-funded projects [11]. For a recent review of nanopore-based methods, see [12]. Specific studies can be found in [13-20].

## 2. Sequencing of whole proteins without proteolysis and/or purification/separation

Most protein sequencing methods, whether at the bulk or at the SM level, are destructive of the sample. Thus a sample protein may be proteolyzed with an enzyme into its constituent peptides, which are then sequenced separately or together. Other methods may sequentially cleave the terminal residue and identify it. Sequencing may be done of a peptide in the protein rather than of the whole protein, or of a subsequence of the protein. Following this the protein may be identified by comparison with sequences stored in a database. Ideally it must be possible to sequence a protein *de novo* without any reference to a database or without prior knowledge of the protein. As a result MS and antibody-based methods are strictly not *de novo*.

While non-destructive methods are preferred the number of known *de novo* whole protein sequencing methods (in any stage of development) is small. From this perspective nanopore-based methods are more likely to be successful as the protein can be translocated through the pore without proteolysis. Such a sequencing method has some seemingly conflicting requirements:

1. It must be able to translocate a protein molecule through the nanopore unidirectionally, from say the *cis* chamber to *trans*, in an e-cell;
2. The translocation must be at a uniform rate so that transport properties can be modeled/computed/measured predictably;
3. Successive residues while inside the pore or when exiting the pore can be distinguished from one another; this usually requires high-bandwidth detectors, which in turn means very high signal-to-noise ratios. Protein sequencing often assumes that the assay is pure, that is, the analyzed sample is made up of copies of a single protein. In practice a sample is more often than not a mixture of different proteins, as it is often hard to separate the component proteins in a mixture. This is especially true when the sample comes from single cells, where some proteins may occur with such low copy numbers that the purification levels required are generally out of reach with currently available methods. One can now add the following fourth desirable property for nanopore-based protein sequencing:
4. The method must be able to sequence proteins in a mixture without purification and/or separation. One way to identify a monomer in a polymer is to attach a label of some kind, such as a fluorescent dye [21], and use a method like Total Internal Reflection Fluorescence to detect the dye and thence the monomer. Labeling and label detection tend to add labor-intensive stages to the workflow. One can now add a fifth desirable property (regardless of the method used):
5. The method is preferably label-free. During the last several years a large number of nanopore-based protein sequencing methods have been reported; none of them appears to resolve all five problems. The study reported in [22] comes closest. While it succeeds with 1 and 2 it is yet to be applied to full-fledged sequencing. The report shows that by adding guanidium chloride to the solution and using some additional artifices a whole protein is unfolded, drawn to a 5 nm biological nanopore, and translocated unidirectionally from *cis* to *trans* at a roughly constant rate, with residues exiting the pore into *trans* at a rate of about one per μs.

### The present study

The present report builds on [22] and uses a method similar to one used earlier in DNA sequencing with nanopores [23] to show that in theory the first four of the five requirements listed above can be satisfied. It is based on attaching conjugate molecules to all residues of a given AA_i_, 1 ≤ i ≤ 20, to selectively increase the blockade current level. (It requires an experimentally distinguishable extension to a target AA type to detect it, so it is intrinsically in violation of requirement 5.) If the whole protein is translocated through the pore as in [22], then with low-frequency low-precision volume measurements (where residue volume acts as a proxy for the blockade level) the partial sequence consisting of AA_i_ can be obtained. By repeating this with all 20 standard AAs, the full sequence can be constructed *de novo*, thus no prior knowledge of the protein or database comparisons are needed. Computational results are given for a sample protein from the human proteome (Uniprot id UP000005640_9606). The model is extended to sequencing of unknown whole proteins in a mixture containing M protein molecules without their separation or purification; copy counts are obtained as a byproduct. The complexity of the sequence extraction algorithm given here is O(N) for a single protein with N residues and O(MN) for a mixture. The potential for translating the procedure into practice either in the lab or in the field is discussed; related implementation issues are addressed.

## 3. Whole protein sequencing with a nanopore from subsequence volumes: A computational model

While events corresponding to residues exiting from the pore in succession can be observed with a time resolution of 1 μs as described in [22], the residue itself cannot be identified exactly or even approximately (such as, for example, belonging to a subset of the 20 AAs [24]). This is because the exclusion volume resolution available is currently limited, the best known to date is ∼70 nm^3^ [24]. Table 1 shows the volumes (in nm^3^) of the 20 AAs. Many of the values in the table are close to each other, requiring very high measurement precision to discriminate among the 20 AAs based on single residue volumes. For example the volumes of M (methionine) and I (isoleucine) differ by 0.9 nm^3^; at present distinguishing these two AAs is impractical if not impossible.

**Table 1.** Volumes (nm^3^) of the 20 standard AAs, data from [25].

| AA | G | A | S | C | D | T | N | P | V | E |
| --- | --- | --- | --- | --- | --- | --- | --- | --- | --- | --- |
| Mean vol | 59.9 | 87.8 | 91.7 | 105.4 | 115.4 | 118.3 | 120.1 | 123.3 | 138.8 | 140.9 |
| AA | Q | H | M | I | L | K | R | F | Y | W |
| Mean vol | 145.1 | 156.3 | 165.2 | 166.1 | 168 | 172.7 | 188.2 | 189.7 | 191.2 | 227.9 |

To get around this problem residue identification can be based on measurement of subsequence volumes. (In [26] such an approach was suggested for protein identification; it was shown in theory that over 90% of the proteins in the proteome of *H. pylori* (UniProt id UP000000210) can be identified with subsequences of length 4 to 8.) In the present study a residue exiting from the pore is associated with the volume excluded by it and its K-1 successor residues during the exit interval of 1 μs. If now a conjugate is added to all instances of AA_i_ in the protein sequence, the \volume of the subsequence inside the pore will be biased near the positions of occurrence of AA_i_ in the sequence. From the biased volume sequence a partial sequence containing only AA_i_ can be obtained. By repeating this procedure with all 20 AAs and merging the 20 partial sequences the whole sequence can be computed.

Table 2 shows the modified volume table that would be used by the sequencing procedure if a conjugate of volume ΔV is attached to all instances of AA alanine (A). (The value of ΔV used below is 1400-2000 nm^3^ This choice for the conjugate volume is discussed below.)

**Table 2.**
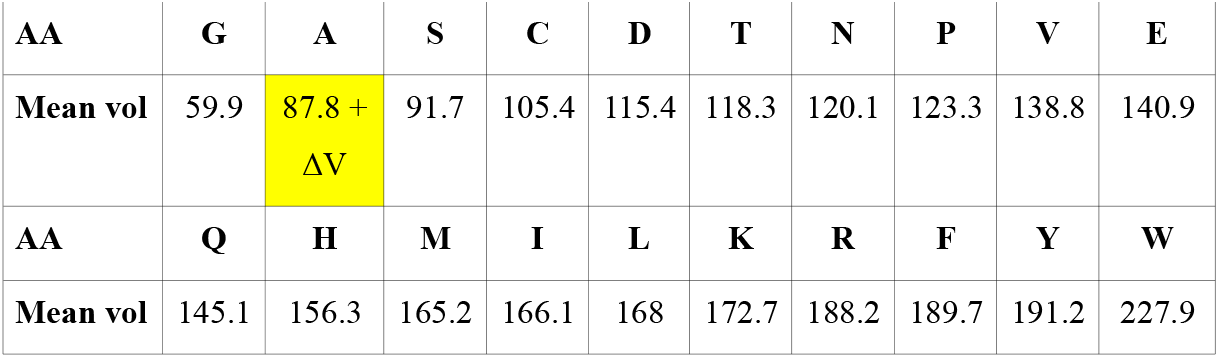
Modified version of Table 1 with volume of A augmented by attached tag of volume ΔV nm^3^.

In the following paragraphs the above notions are used to formally develop a sequencing procedure for whole proteins.

### Assumptions

1. With an e-cell in which analytes (that is, sample proteins) are loaded into the *cis* chamber and translocate to the *trans* chamber, translocation signals can be measured with a time resolution of 1 μs [22]
2. A protein is in a stretched state as it translocates through the pore unidirectionally from *cis* to *trans*. In [22] the pore used is the biological pore alpha Hemolysin (AHL), which has a length of L = 5 nm. With such a pore there are 10 residues inside the pore at any time. A profile dimension per residue (its length along the pore axis) of 0.5 nm is assumed for all residues. (The residue to residue distance along the backbone of a linear stretched protein is roughly the same for all residue pairs; AA volume differences are largely in the side chain.) With this assumption there are K = 2L = 10 residues inside the pore at any time.
3. Successive residues exit at a rate of 1/μs at the *trans* end of the pore into *trans* without regressing.
4. Analyte volume is used as a proxy for the blockade current level. Current blockades in a nanopore occur due to the analyte physically excluding current carriers (ions in an electrolyte, typically K^+^ and Cl^-^ in a solution containing KCl) in the volume occupied by the analyte while it translocates through the pore. This exclusion volume is assumed to be proportional to the drop in the current that would normally flow in the absence of an analyte.
5. The resolution with which blockade exclusion volumes are measured is ∼70 nm^3^. (This level of precision was used in [24] to distinguish among four subsets of the AAs labeled S(mall), M(edium), I(ntermediate), and L(arge) according to their volumes.)

### Notation

k = index of residue at the *trans* end of the pore

Δt = time duration of exit of *trans* end residue into *trans*

K = number of residues in protein segment inside pore (= 10 = 2L for a pore of length L = 5 nm)

p_i_ = i-th residue in protein

Vol_i_ = volume of p_i_ (see Table 1)

p_k_ p_k+1_ … p_k+K-1_ = protein segment inside pore (p_k_ = residue at trans end of pore)

psVol_k,K_ = total volume of p_k_ p_k+1_ … p_k+K-1_ (there are K residues in the pore at any time)

cpsVol_k,K_ = total volume of p_k_ p_k+1_ … p_k+K-1_ with AA_i_ occurrences conjugated

T_k,m_= translocation time of subsequence of length k starting at m

V_step_ = volume precision = 70 nm^3^

dV_x_ = digitized volume of V_x_ = ⌊V_x_/V_step_⌋

Consider a protein P with N residues given by

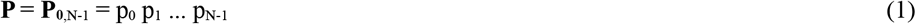

A segment with K = 10 residues from the k-th to the (k+K-1)-th that are resident in the pore is given by

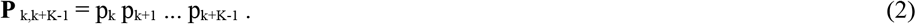

Now consider a whole protein translocating from *cis* to *trans* through the pore. The time taken for the segment inside the pore **P** _k,k+M-1_ = p_k_ p_k+1_ … p_k+M-1_, k=0,1,…N-K+1 to translocate out can be considered in terms of the times taken by each residue as it exits the pore. Since the latter is the same for all residues [22] this time can be written as

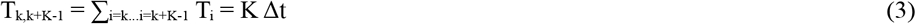

The volume excluded by this segment is the pore sequence volume

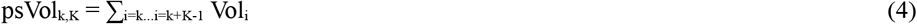

where Vol_i_ is the volume of residue i (Table 1).

Given the limited volume measurement precision of V_step_ (= 70 nm^3^, see above), the measured volume is the digitized pore sequence volume

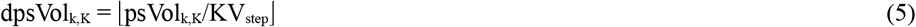

The digitization in Equation 5 may lead to truncation errors, this is taken into account in the sequencing procedure given below.

To obtain the exact identity of a residue from these digitized volumes an artifice analogous to the use of base-specific tags for DNA sequencing with a nanopore [23] is used. In [23] the tag molecules are base-specific and are attached to the corresponding bases in a DNA sequence. The DNA strand is pulled with an enzyme motor attached to the membrane near the entrance to the pore. A tag is removed by a chemical reaction when the base comes near the pore entrance. The released tag falls through the pore causing a current blockade of a unique size. As the strand ratchets through the motor the tags of successive bases are detached and identified when they fall through the pore so that the sequence of blockade levels caused by them unambiguously yields the base sequence of the strand.

In the present case conjugates that add volume ΔV are attached to a given AA type, this pushes up the measured volume of the pore sequence near the conjugated AA, from which the position of occurrence of every residue of the type in the whole sequence is extracted. By applying this step separately to each of the 20 standard AAs the full sequence can be obtained. The value chosen for ΔV is based on the following considerations:

1. With a volume measurement precision of 70 nm^3^ the measured exclusion volume due to the 10 residues resident in the pore at any time is on the order of 700 nm^3^. Digitization imposes a truncation (or roundoff) error of up to 700 nm on the pore sequence volume that is sampled every μs. With a conjugate volume ΔV = 1400 nm^3^ or more there is a minimum change in the signal volume of 700 nm^3^.
2. The unconjugated volume varies somewhat over the full length of the protein. This is further affected by truncation/roundoff errors in digitization.

With residues of a selected AA type conjugated Equations 4 and 5 can be rewritten as the conjugated pore sequence volumes before and after digitization:

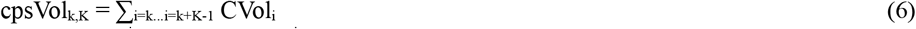

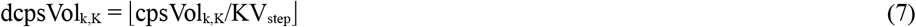

where CVol_i_ is the volume of residue i from Table 2. Equation 8 gives the signal corresponding to the difference between the unconjugated sequence volume and conjugated sequence volume:

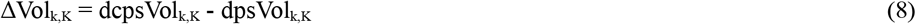

Rewrite Equation 7 as

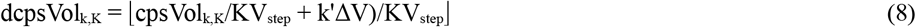

where k’ is the number of occurrences of the conjugated AA_i_ in [k, k+K-1]. If ΔV is set to k’’ KV_step_, where k’’ is a small positive integer (1 or 2), then

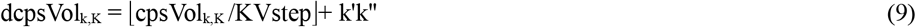

With k’’ = 2, the added volume is 1400 nm^3^ and the digitized pore sequence volume increases by an integer value of 2 for every occurrence of the the conjugated AA_i_ in the pore sequence. Effectively a conjugated AA_i_ at position m adds a digital volume of 2 to the pore volume sequence at positions u in [m-(M-1)+1, m], this added volume is removed when the residue leaves the pore. This effectively adds a rectangle of height 2 to the digitized unconjugated sequence signal; the rectangles may overlap. Sequencing in effect reduces to finding the positions of each rectangle, overlapping or not. The algorithm to do this is given next. It is based on locating the start of the first rectangle, subtracting this rectangle from the difference between the conjugated and unconjugated sampled volume sequences, locating the start of the next rectangle, and repeating this process over the full signal sequence. Finding the start of the first of the remaining rectangles in this process consists of scanning the remaining sequence to locate the rising edge of that rectangle. This procedure returns 20 lists with the positions of occurrence of a conjugated AA, one list per AA. Following this the 20 lists are merged into a single list based on the positions of each AA. Merging can be done with a 20-way sorted merge. The pseudocode for is given below in the two procedures.

**Procedure** FindRectanglesforConjugatedResidues

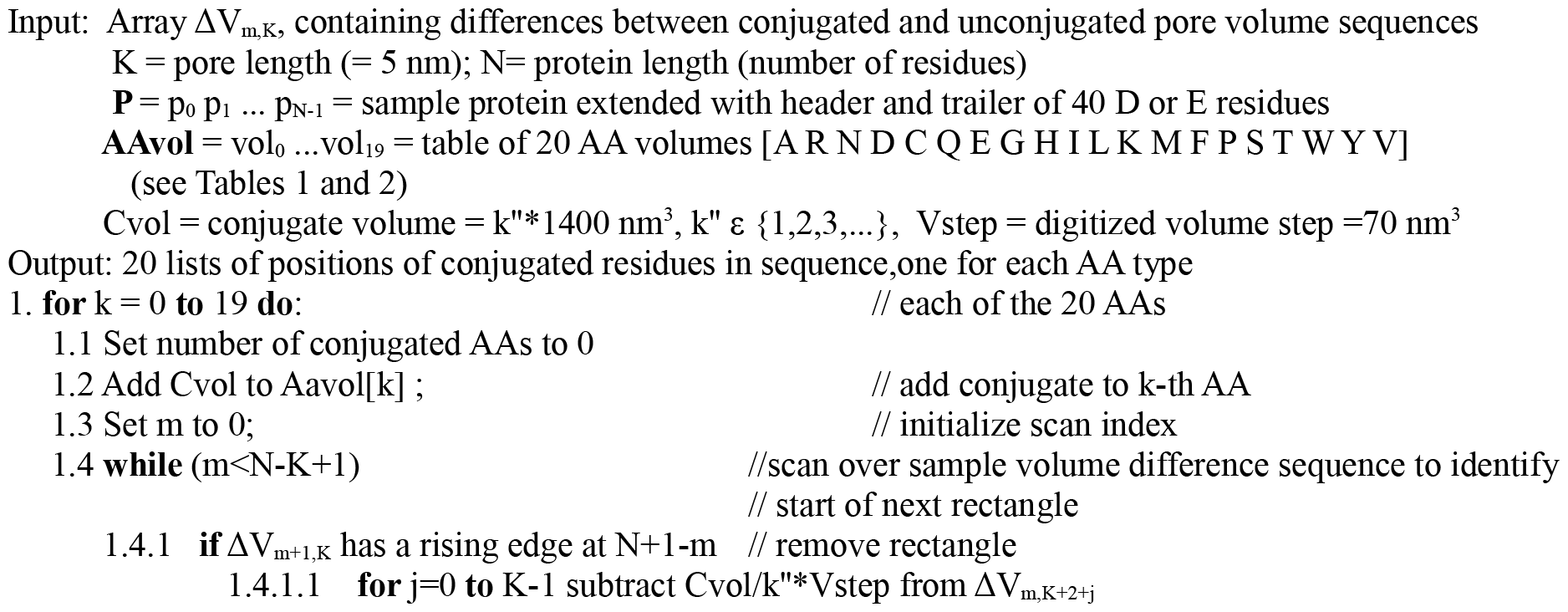
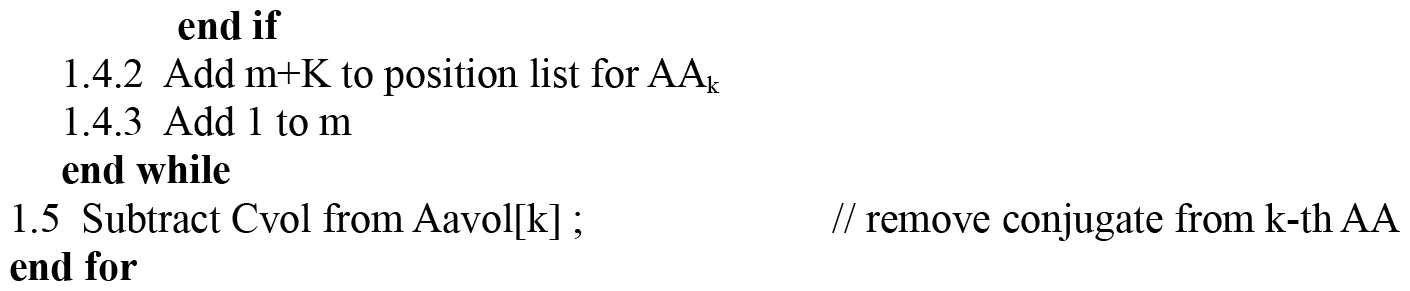

**Procedure** FindProteinSequence

Input: 20 lists of positions of conjugated AA, one per AA Output: Sequence of AAs for protein

1. Call K-WayMergeSort algorithm with K = M = 20 to merge-sort 20 sequences of integers corresponding to positions of AA in conjugated sequence

The above method will give exact sequencing information for all 20 conjugated sequences. Less rigid values can be used for ΔV with somewhat more involved heuristics but the results may contain false positives and false negatives. Thus let ΔV be set to (k’’ + Δ) KV_step_, + where Δ, 0 < Δ < 1. Then Equation 9 becomes

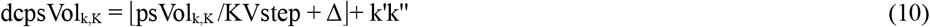

The height of the rectangle resulting from a single conjugated AA is no longer fixed at k’’, it could be k’’ or k’’ +1, which could cause the algorithm given above to produce false positives or false negatives. Better heuristics may be used to reduce the number of false positives and negatives to a negligible level.

### Complexity of computing the sequence of a single protein

In FindRectanglesforConjugatedResidues the while loop moves the scanner m from to near the end of the sequence of length N, recording successively increasing values for the positions of the conjugated AA in the subsequence for A. This a one-way move, there is no backtracking. The resulting complexity is thus of order O(N).

In FindProteinSequence the 20 lists carrying position information for the 20 AAs are all sorted on the position of conjugated AAs. As a result the merge operation is once again a one-way move without any backtracking, the complexity is O(N).

The overall complexity of computing the sequence of a single protein from the 20 lists of conjugated AA positions is thus O(N).

The above development is illustrated in the next section with an example protein and ΔV values of 1400 and 1500.

## 4 Computational results: An example

The whole protein sequencing procedure described above is illustrated with protein 5 (Uniprot id A6NL46, 340 residues) in the human proteome (Uniprot id UP000005640_9606). The top half of Figure 1 shows the protein sequence. The bottom half shows the sequence with a header and trailer of 40 E (glutamic acid) residues each.

**Figure 1.**
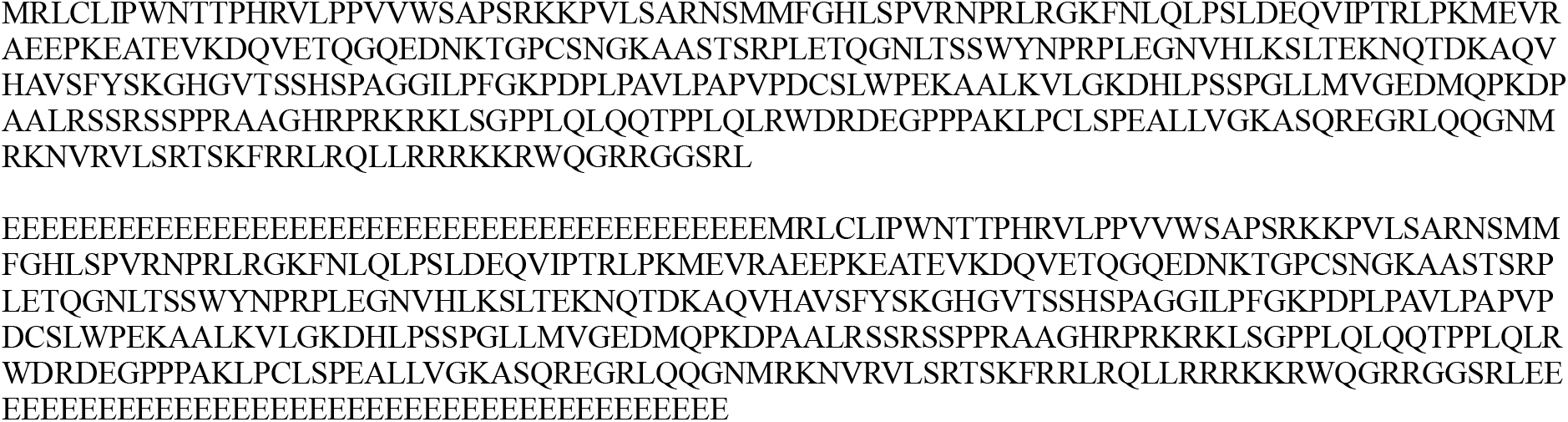
Protein 5, id A6NL46, in human proteome (Uniprot UP000005640_9606)

Figure 2 shows the sequence of residue volumes for the first 345 residues; for space reasons the x axis displays every other residue.

**Figure 2.**
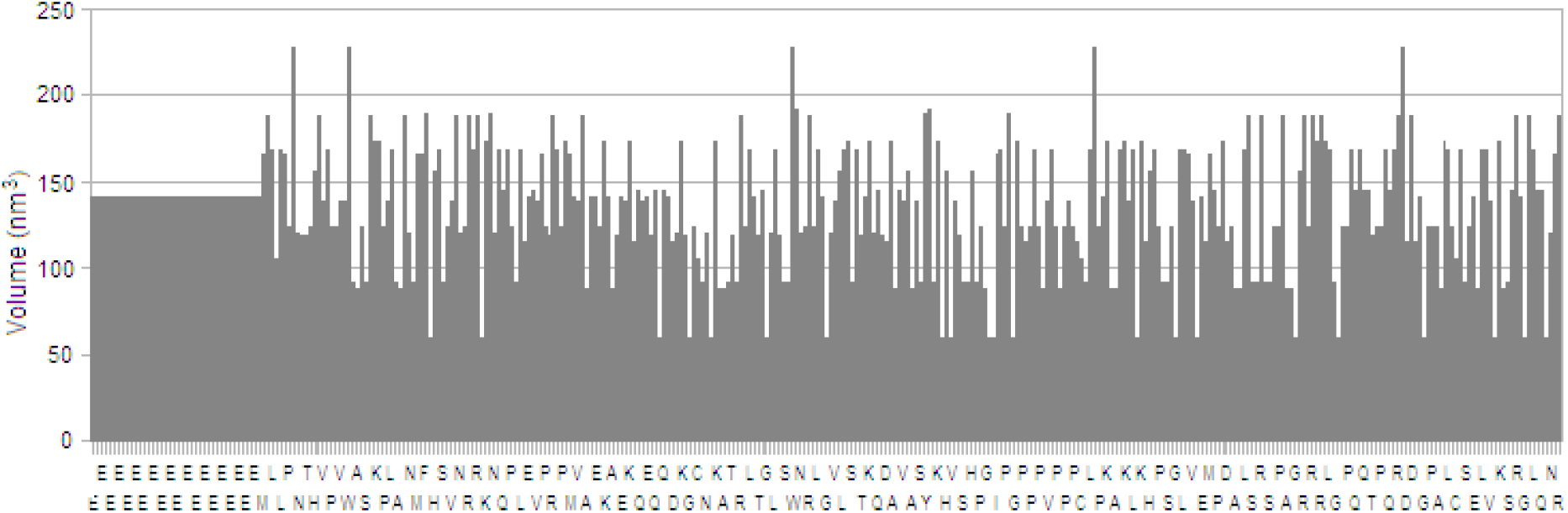
Residue volumes of the first 345 residues in Protein 5, id A6NL46, in human proteome

**Figure 3.**
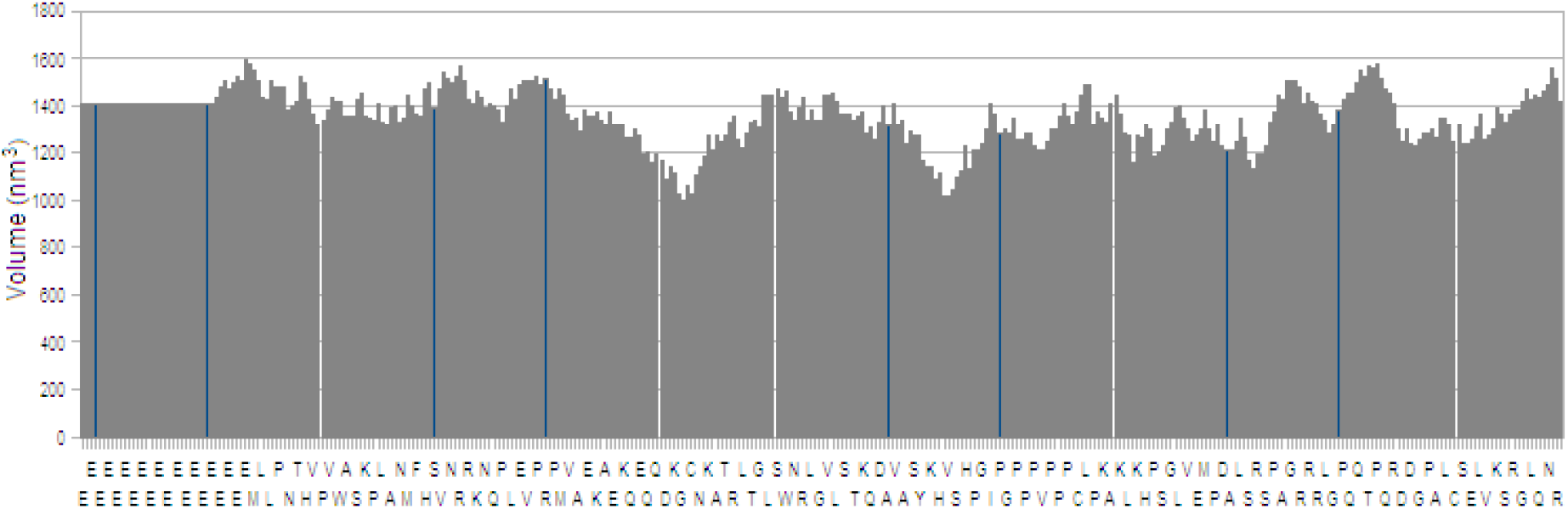
Graph of measured pore sequence volumes along protein sequence (first 345 residues shown)

**Figure 4.**
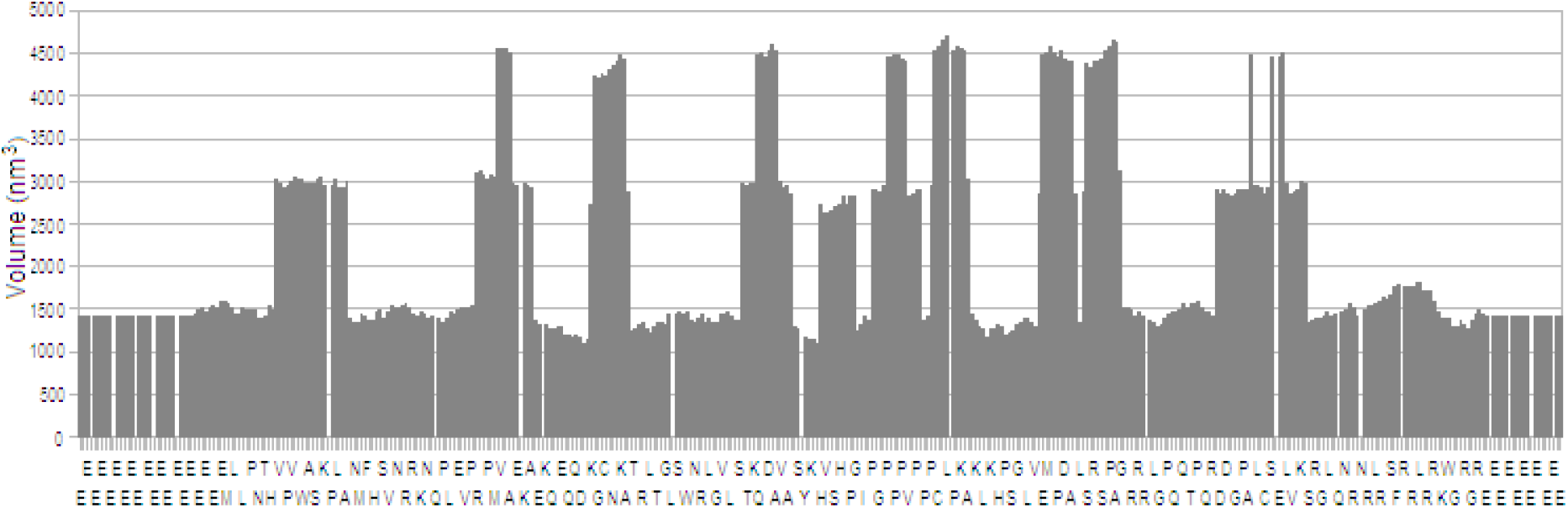
Graph of measured pore sequence volumes along protein sequence with alanines conjugated (first 345 residues in sequence are shown). There are 20 alanines in the full sequence, those to the right of position 345 are not shown.

**Figure 5.**
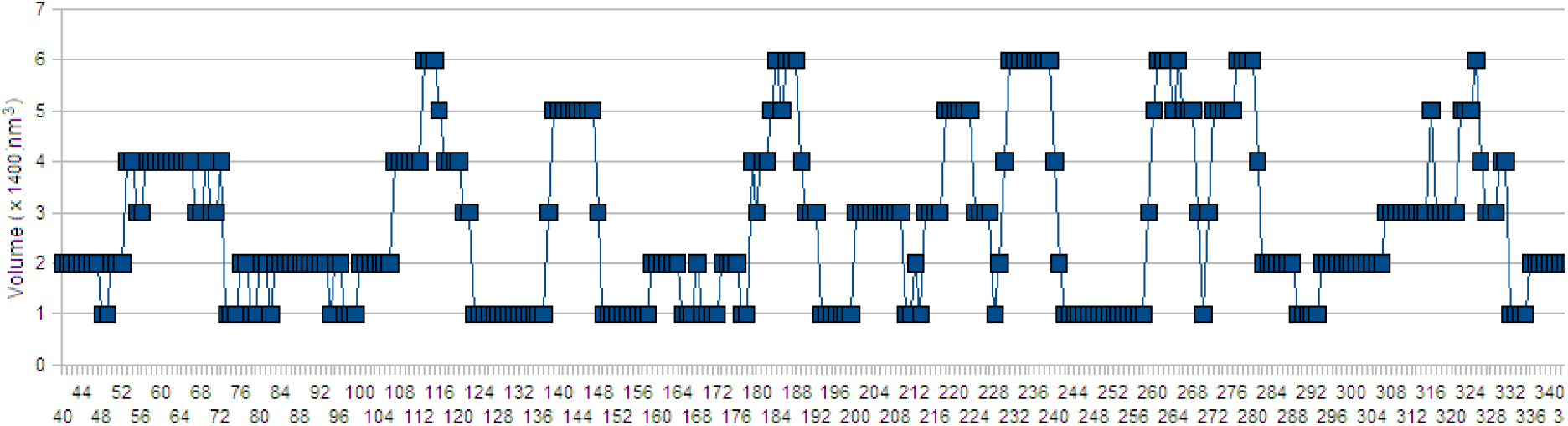
Graph of digitized conjugated sequence volumes in Figure 4 with conjugate volume of 1400 nm^3^

**Figure 6.**
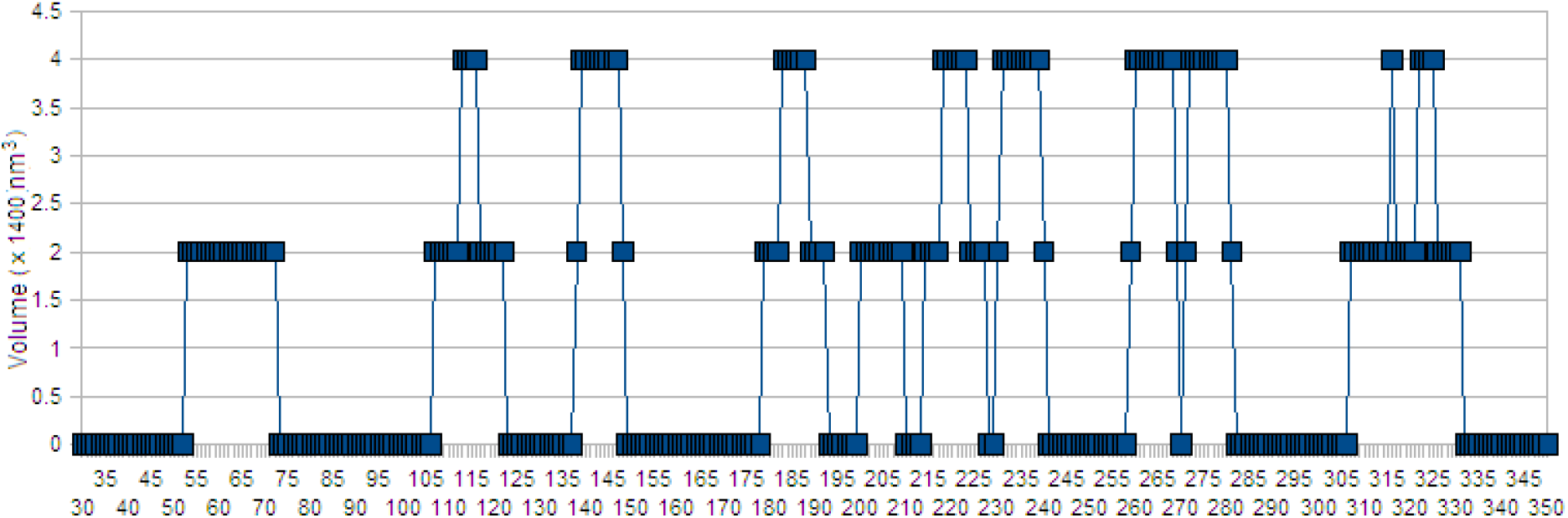
Graph of difference between digitized unconjugated pore sequence volumes in Figure 3 and digitized conjugated sequence volume with conjugate volume of 1400 nm^3^ from Figure 5

**Figure 7.**
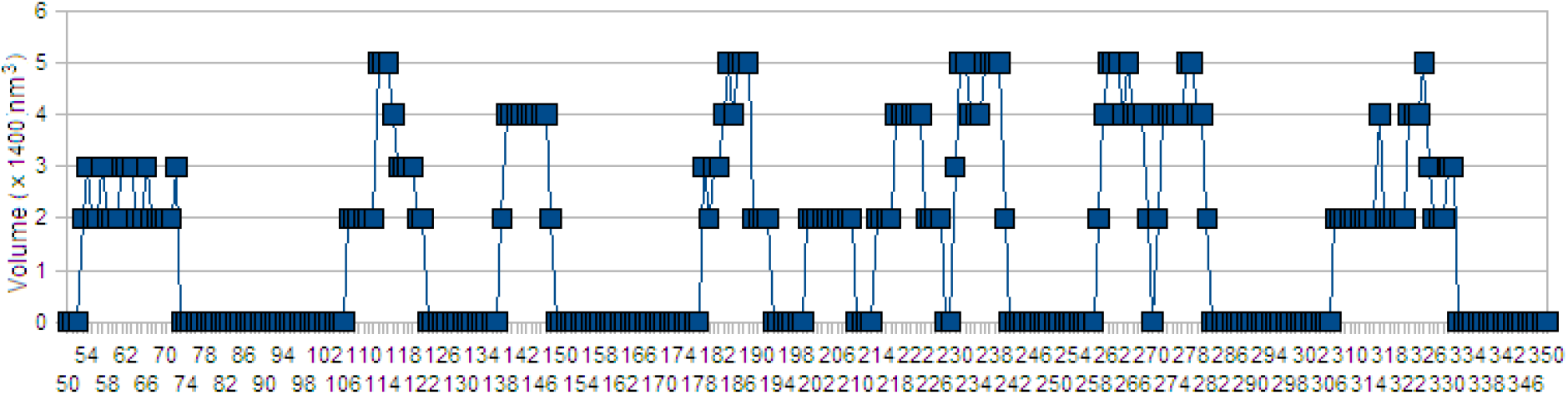
Similar to Figure 6 but with conjugate volume of 1500 nm^3^

Figure 8 shows the actual and computed sequences with ΔV = 1400 nm^3^. The match is exact for reasons discussed above, see Section 3. Figure 9 shows the actual and computed sequences with ΔV = 1500 nm^3^. There are a number of mismatches, resulting in false positives and false negatives. As noted earlier, these mismatches can be traced to to the truncation error that occurs in Equations 7-9.

**Figure 8.**
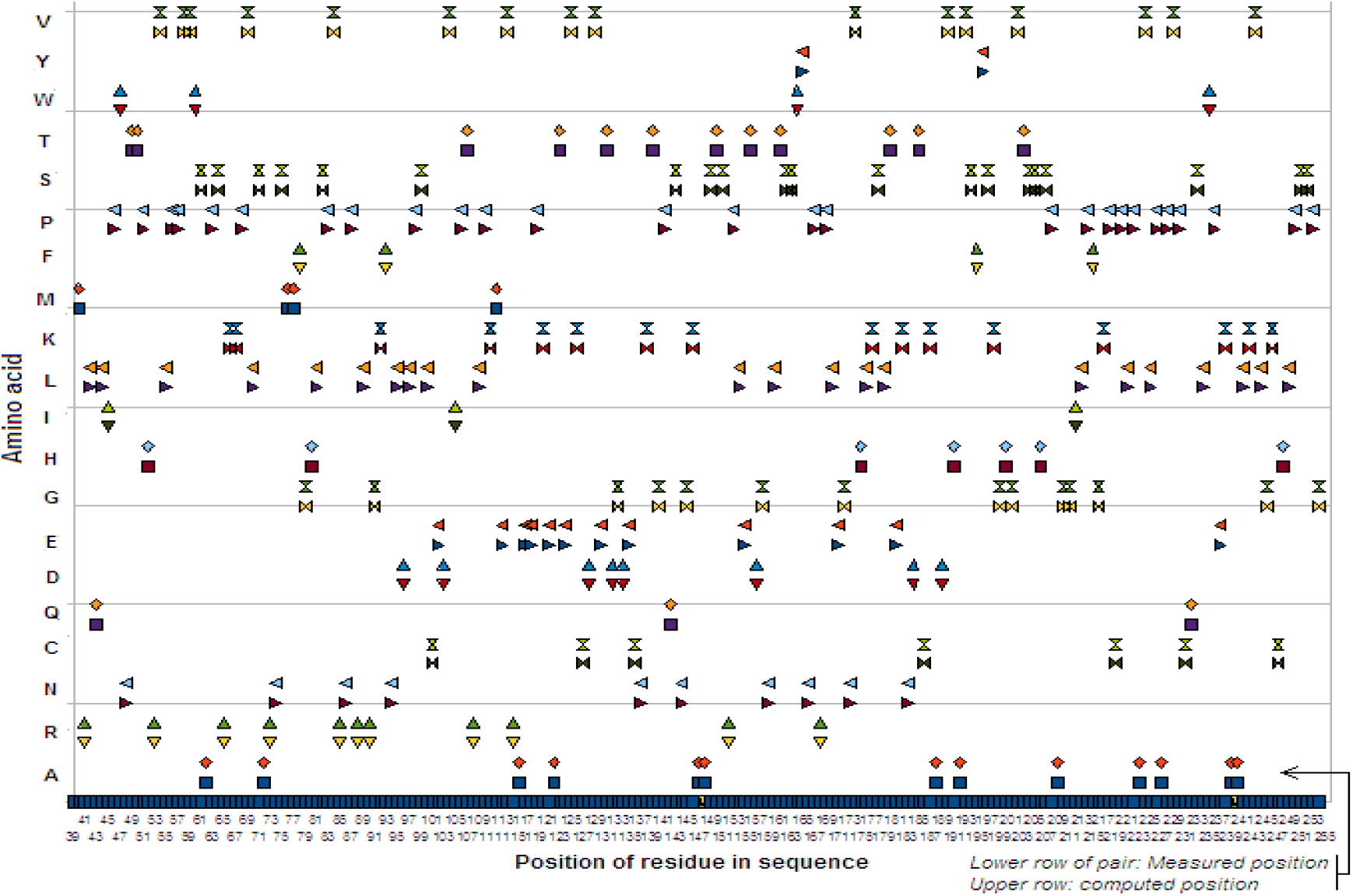
Computed sequence matches exactly with actual sequence when ΔV = 1400

**Figure 9.**
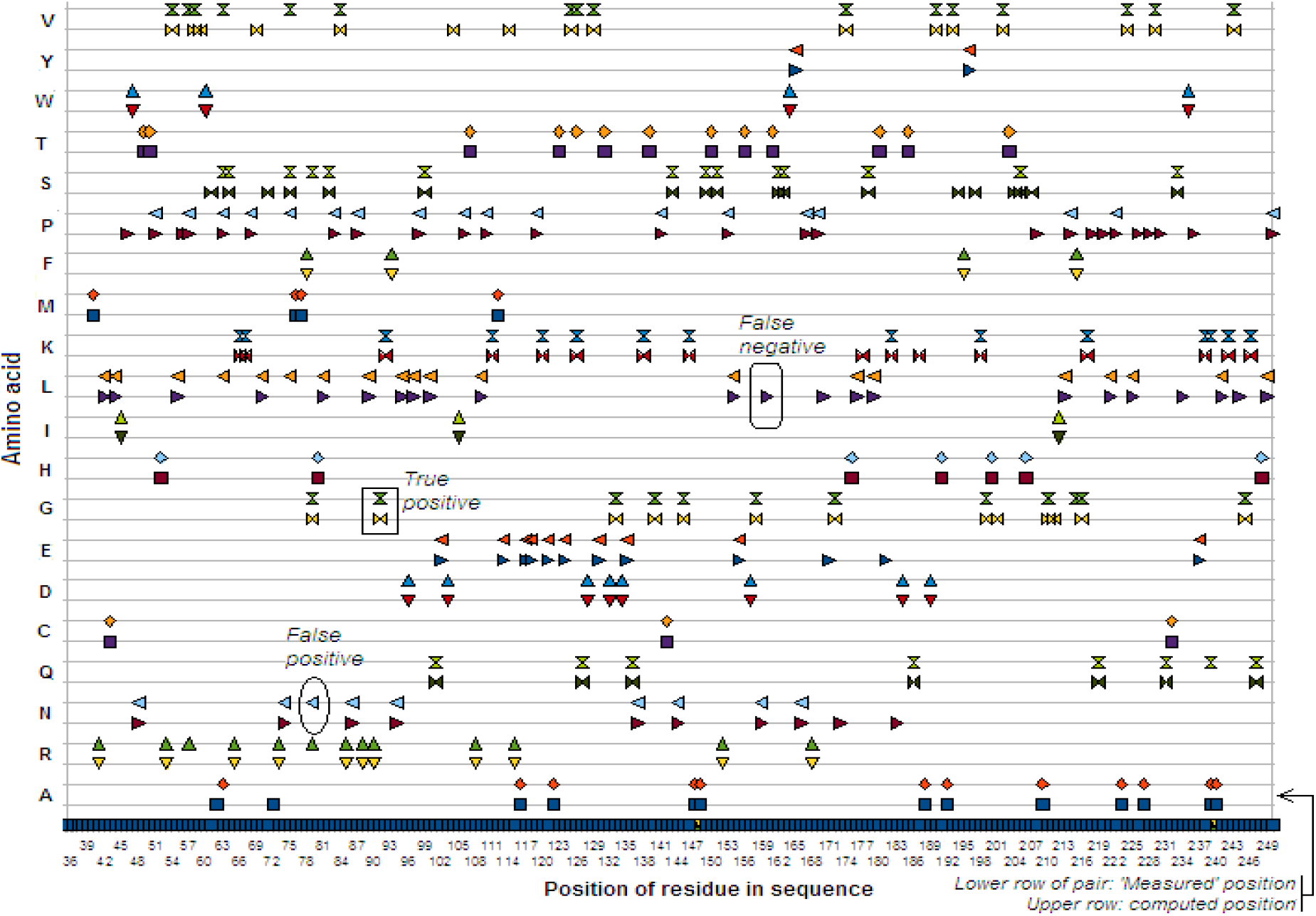
Computed sequence matches and actual sequence when ΔV = 1500. Examples of true match, false positive, and false negative are shown in enclosing boxes of different shapes.

Tables 3A and 3B show lists of actual and detected positions of all 20 AAs in the protein sequence when ΔV = 1500 nm^3^. At the bottom of each table the number of false positives and of false negatives for each AA is given. Table 4 collects these numbers in a single place for convenience and also includes the total number for errors.

**Table 3A.**
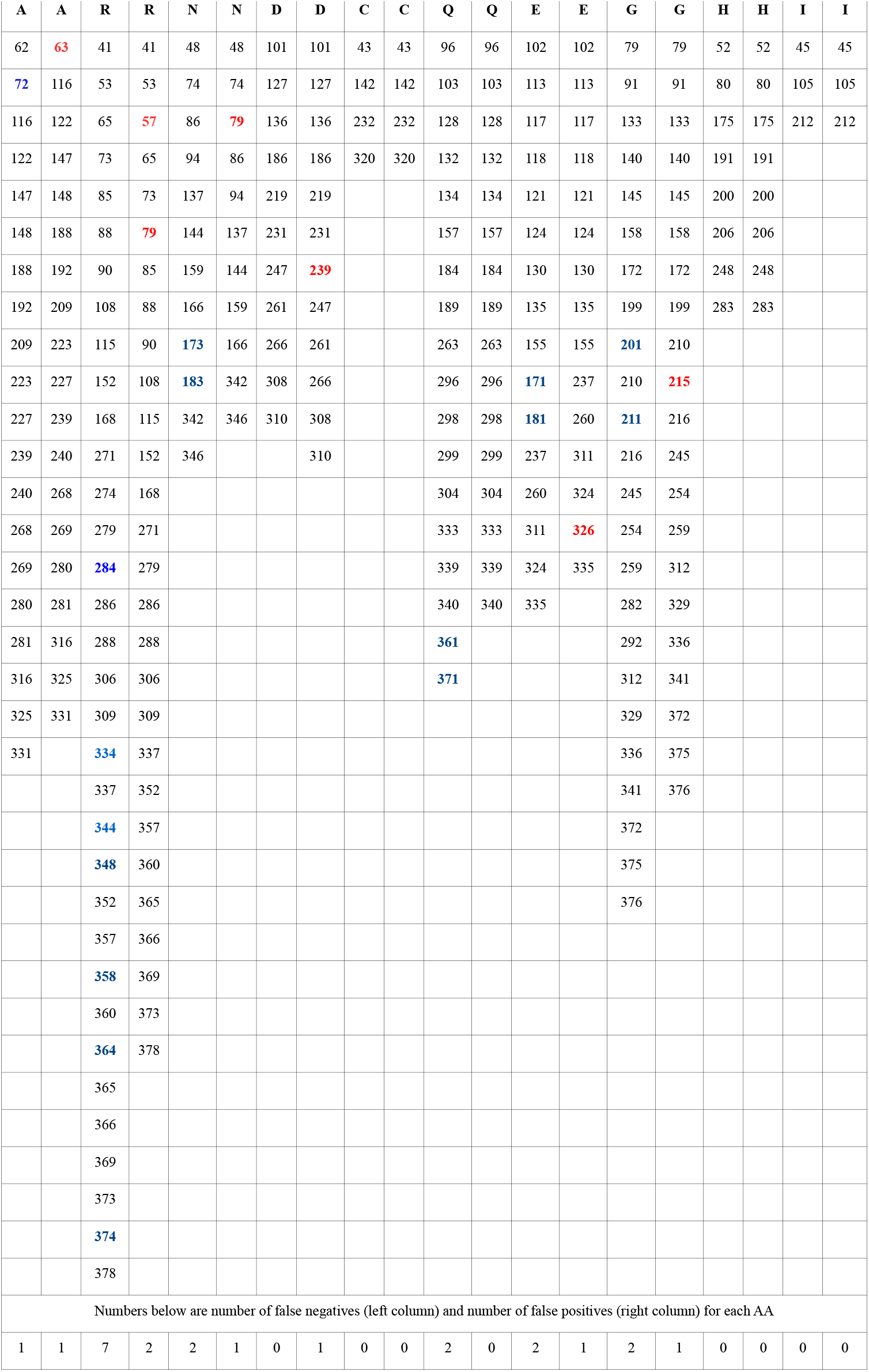
Positions of AAs (left column of AA) in actual sequence and in computed sequence (right column) with ΔV = 1500. Blue: false negative, Red: False positive.

**Table 3B.** Positions of AAs (left column of AA) in actual sequence and in computed sequence (right column) with ΔV = 1500. Blue: false negative, Red: False positive.

| L | L | K | K | M | M | F | F | P | P | S | S | T | T | W | W | Y | Y | V | V |
| --- | --- | --- | --- | --- | --- | --- | --- | --- | --- | --- | --- | --- | --- | --- | --- | --- | --- | --- | --- |
| 42 | 42 | 66 | 66 | 40 | 40 | 78 | 78 | 46 | 51 | 61 | 63 | 49 | 49 | 47 | 47 | 165 | 165 | 54 | 54 |
| 44 | 44 | 67 | 67 | 76 | 76 | 93 | 93 | 51 | 57 | 64 | 64 | 50 | 50 | 60 | 60 | 196 | 196 | 58 | 57 |
| 55 | 55 | 92 | 92 | 77 | 77 | 195 | 195 | 56 | 63 | 71 | 75 | 107 | 107 | 164 | 164 |  |  | 59 | 58 |
| 70 | 63 | 111 | 111 | 112 | 112 | 215 | 215 | 57 | 68 | 75 | 79 | 123 | 123 | 235 | 235 |  |  | 69 | 63 |
| 81 | 70 | 120 | 120 | 257 | 257 | 356 | 356 | 63 | 75 | 82 | 82 | 131 | 126 | 307 | 307 |  |  | 84 | 75 |
| 89 | 75 | 126 | 126 | 262 | 262 |  |  | 68 | 83 | 99 | 99 | 139 | 131 | 370 | 370 |  |  | 104 | 84 |
| 95 | 81 | 138 | 138 | 343 | 343 |  |  | 83 | 87 | 143 | 143 | 150 | 139 |  |  |  |  | 114 | 125 |
| 97 | 89 | 146 | 146 |  |  |  |  | 87 | 98 | 149 | 149 | 156 | 150 |  |  |  |  | 125 | 126 |
| 100 | 95 | 177 | 182 |  |  |  |  | 98 | 106 | 151 | 151 | 161 | 156 |  |  |  |  | 129 | 129 |
| 109 | 97 | 182 | 198 |  |  |  |  | 106 | 110 | 162 | 162 | 180 | 161 |  |  |  |  | 174 | 174 |
| 154 | 100 | 187 | 217 |  |  |  |  | 110 | 119 | 163 | 163 | 185 | 180 |  |  |  |  | 190 | 190 |
| 160 | 109 | 198 | 238 |  |  |  |  | 119 | 141 | 178 | 178 | 203 | 185 |  |  |  |  | 193 | 193 |
| 170 | 154 | 217 | 239 |  |  |  |  | 141 | 153 | 194 | 205 | 300 | 203 |  |  |  |  | 202 | 202 |
| 176 | 176 | 238 | 242 |  |  |  |  | 153 | 167 | 197 | 233 | 353 | 300 |  |  |  |  | 224 | 224 |
| 179 | 179 | 242 | 246 |  |  |  |  | 167 | 169 | 204 | 251 |  | 353 |  |  |  |  | 229 | 229 |
| 213 | 213 | 246 | 265 |  |  |  |  | 169 | 214 | 205 | 252 |  |  |  |  |  |  | 243 | 243 |
| 221 | 221 | 265 | 287 |  |  |  |  | 208 | 222 | 207 | 272 |  |  |  |  |  |  | 258 | 258 |
| 225 | 225 | 287 | 289 |  |  |  |  | 214 | 250 | 233 | 273 |  |  |  |  |  |  | 328 | 328 |
| 234 | 241 | 289 | 317 |  |  |  |  | 218 | 253 | 251 | 275 |  |  |  |  |  |  | 347 | 347 |
| 241 | 249 | 317 | 326 |  |  |  |  | 220 | 264 | 252 | 276 |  |  |  |  |  |  | 349 | 349 |
| 244 | 255 | 330 | 330 |  |  |  |  | 222 | 278 | 272 | 291 |  |  |  |  |  |  |  |  |
| 249 | 256 | 345 | 367 |  |  |  |  | 226 | 285 | 273 | 326 |  |  |  |  |  |  |  |  |
| 255 | 270 | 355 | 368 |  |  |  |  | 228 | 293 | 275 | 351 |  |  |  |  |  |  |  |  |
| 256 | 290 | 367 |  |  |  |  |  | 230 | 294 | 276 | 354 |  |  |  |  |  |  |  |  |
| 270 | 297 | 368 |  |  |  |  |  | 236 | 301 | 291 | 377 |  |  |  |  |  |  |  |  |
| 290 | 303 |  |  |  |  |  |  | 250 | 302 | 322 |  |  |  |  |  |  |  |  |  |
| 295 | 318 |  |  |  |  |  |  | 253 | 314 | 332 |  |  |  |  |  |  |  |  |  |
| 297 | 321 |  |  |  |  |  |  | 264 | 315 | 351 |  |  |  |  |  |  |  |  |  |
| 303 | 326 |  |  |  |  |  |  | 267 | 319 | 354 |  |  |  |  |  |  |  |  |  |
| 305 | 327 |  |  |  |  |  |  | 277 |  | 377 |  |  |  |  |  |  |  |  |  |
| 318 | 338 |  |  |  |  |  |  | 278 |  |  |  |  |  |  |  |  |  |  |  |
| 321 | 350 |  |  |  |  |  |  | 285 |  |  |  |  |  |  |  |  |  |  |  |
| 326 | 359 |  |  |  |  |  |  | 293 |  |  |  |  |  |  |  |  |  |  |  |
| 327 | 362 |  |  |  |  |  |  | 294 |  |  |  |  |  |  |  |  |  |  |  |
| 338 | 363 |  |  |  |  |  |  | 301 |  |  |  |  |  |  |  |  |  |  |  |
| 350 | 379 |  |  |  |  |  |  | 302 |  |  |  |  |  |  |  |  |  |  |  |
| 359 |  |  |  |  |  |  |  | 313 |  |  |  |  |  |  |  |  |  |  |  |
| 362 |  |  |  |  |  |  |  | 314 |  |  |  |  |  |  |  |  |  |  |  |
| 363 |  |  |  |  |  |  |  | 315 |  |  |  |  |  |  |  |  |  |  |  |
| 379 |  |  |  |  |  |  |  | 319 |  |  |  |  |  |  |  |  |  |  |  |
|  |  |  |  |  |  |  |  | 323 |  |  |  |  |  |  |  |  |  |  |  |
| Numbers below are number of false negatives (left column) and number of false positives (right column) for each AA |  |  |  |  |  |  |  |  |  |  |  |  |  |  |  |  |  |  |  |
| 5 | 1 | 4 | 2 | 0 | 0 | 0 | 0 | 11 | 3 | 7 | 2 | 0 | 1 | 0 | 0 | 0 | 4 | 0 | 2 |

**Table 4.** False positives and false negatives in computed sequence for protein 5 with ΔV = 1500 nm^3^

| AA | A | R | N | D | C | Q | E | G | H | I | L | K | M | F | P | S | T | W | Y | V | Total |
| --- | --- | --- | --- | --- | --- | --- | --- | --- | --- | --- | --- | --- | --- | --- | --- | --- | --- | --- | --- | --- | --- |
| No of false positives | 1 | 7 | 2 | 0 | 0 | 2 | 2 | 2 | 0 | 0 | 5 | 4 | 0 | 0 | 11 | 7 | 0 | 0 | 0 | 4 | 47 |
| No of false negatives | 1 | 2 | 1 | 1 | 0 | 0 | 1 | 1 | 0 | 0 | 1 | 2 | 0 | 0 | 3 | 2 | 1 | 0 | 0 | 2 | 18 |

## 5. Sequencing and quantification of proteins in a mixture

Consider a mixture of M protein molecules P_1_, P_2_, …, P_M_. The output of the sequencing procedure described in the previous section when applied to the mixture consists of 20M partial sequences. For each of the M molecules there are 20 conjugated sequences of length N, one per AA_k_, 1 ≤ k ≤ 20. Sequencing consists of finding the set of conjugated sequences corresponding to the same molecule. One way to do this is to map the lists of AA-specific residue positions for each conjugated sequence to an undirected 20-partite graph G_M_ with 20M vertexes, one list per matrix entry. Two vertexes in G_M_ are adjacent if and only if their lists have no common members. All vertexes that are adjacent to each other correspond to conjugated sequences for a single molecule; such vertexes form a clique. Thus G_M_ contains M disjoint cliques of degree 20. Sequencing of the M molecules then reduces to identifying these cliques. The clique identification problem is computationally hard [27]; an alternative approach that is nominally in polynomial time is described next.

The algorithm to use is a straightforward one that works with an M × 20 matrix that has M rows for the M protein molecules (in some unknown order) and 20 columns corresponding to AA-specific subsequences obtained by measurement for each AA type. Every column is independent of the other columns, the column entries may be in some random order of the M molecules as the measurement process has no knowledge of the identity of the molecule. Sequencing of a protein molecule consists of bringing together one entry from each of the 20 columns equal to the subsequences for the AA corresponding to that column. This can be done by finding M disjoint paths through the M columns such that each of the M sets of subsequences along the respective paths forms a full protein sequence when merged. The algorithm to do this assembly, Procedure *AssembleSequencesFromConjugateSequenceMatrix*, is given next. It is followed by an algorithm for quantification, Procedure *QuantifySampleFromMProteinSequences*.

**Procedure** *AssembleSequencesFromConjugateSequenceMatrix*

Input: Conjugate sequence matrix CSM[M × 20 ] (see Appendix for example with 6 proteins from the human proteome (Uniprot id UP000005640_9606; protein nos. 4, 5, 7, 8, 10, and 16))

Output: M sequences of residues for M protein molecules

Use depth-first search to traverse over a path of length 20: [i_1_, j_1_] [i_2_, j_2_]…[i_20_, j_20_], 1 ≤ i_1_, i_2_, … i_20_ ≤ M, 1 ≤ j_1_, j_2_, … j_20_ ≤ 20, with all j’s distinct, through the matrix CSM. If the assembled set of positions from each matrix entry along a path is the integer sequence 0,…,N-1 the sequence of AAs along the path corresponds to a protein. There should be exactly M such paths corresponding to the M proteins in the sample.

When there are multiple copies of a protein in the sample, the value of M can be reduced considerably before proceeding to assemble the M protein sequences with *AssembleSequencesFromConjugateSequenceMatrix*. With duplicates sequence determination is markedly easier. This is because if there are d duplicates of a protein there will be d identical entries in each of the 20 columns. These can easily be checked for intersections across the columns, following which assembly of the sequence can be done efficiently. Thereafter these duplicates can be removed from the CSM matrix, the remaining entries in the CSM matrix then correspond to single-copy proteins. d can be as small as 2. Thus if there are 2 or more copies of a protein in the sample, regardless of how many total molecules there are in the sample, its sequence can be obtained quickly. This reduced CSM matrix contains the protein sequences of M single-copy proteins.

Quantification of the sample can be done by sorting the M protein sequences on the sequence strings and counting duplicates down the sorted list.

**Procedure** *QuantifySampleFromMProteinSequences*

Input: M protein sequences output by *AssembleSequencesFromConjugateSequenceMatrix*

Output: M’ distinct protein sequences p_1_1_…p_1_k1_, p_2_1_…p_2_k2_,, …, p_M’_1_…p_M’_kM’_ and their copy numbers q_1_,…,q_M’_ with q_1_ + … + q_M’_ = M

Step 1. Sort the protein sequences on the strings p_1_x_…p_1_y ._

Step 2. In the sorted list count the number of duplicates down the string and output copy count for protein

### Complexity of mixture sequencing and quantification

1. In Procedure *AssembleSequencesFromConjugateSequenceMatrix* there are M^20^ distinct paths of length 20 from the first column to the 20^th^ column of CSM[M][20]. In the worst case all of them may be examined, so the worst-case behavior is O(M^20^). This is nominally in polynomial time but is not practical for values of M exceeding 5 or 6. Notice that once a sequence has been determined the cells in CSM from which the sequence is generated can be removed, this effectively reduces M by 1 after every sequence determination. Thus the number of rows in CSM goes down in M-1 cycles from M to 1. As a result the complexity is somewhat less than O(M^20^).
2. In Procedure *QuantifySampleFromMProteinSequences* the sorting step has complexity O(M log M). The following collation step (Step 2) walks through the sorted list counting the number of entries in each block containing identical strings. This can be done in O(M) time.

Following the removal of proteins with duplicates assume that the reduced CSM is of order M × 20 and contains the conjugate sequence lists for single copy proteins. The mixture sequencing method given here needs to use a sample size with only a small number of single protein molecules. To this end an assay sample can be subdivided into subsamples with M molecules, where M is in a computationally practical range. When the dynamic range of a sample is very large (as is true with single cell assays in which the sample can contain millions of copies of some proteins and very few, possibly in the single digit, of others) suitable statistical sampling methods may be used to ensure that the (statistical) sample is unbiased and the assay sample includes single copy proteins with high probability.

## 6 Discussion

Assuming that conjugate molecules specific to each AA can be designed, it would appear that the technology needed to translate the proposed method into practice is available. A number of implementation and procedural issues are considered next.

1. The proposed procedure may be implemented in the bulk if samples of sufficient size are available.
2. If sufficient quantities of sample molecules are not available the procedure can be implemented serially in 20 cycles, one per AA. Each cycle would then consist of the following sequence of steps: 1. Attach conjugates for AA_i_ to every protein molecule in the sample; 2. Sequence the conjugated sample mixture; 3. Detach conjugates and wash away conjugates. If a sufficient quantity of the sample is available then 20 channels, one per AA type, can be used to obtain conjugated sequences with that AA. Following this the assembly of sequences can be done as in Section 5.
3. With a parallel array of 10000 nanopores, a conservative time estimate of 1 hour for conjugation and 1 hour for detaching and washing, and negligible data processing time, 10000 samples with M molecules in each can be sequenced and quantified in ∼2 hours.
4. In [22] a header and trailer of 20-30 charged D residues is used to draw the protein into the pore and encourage unidirectional translocation. Incidentally this artifice can also be used to determine the entry orientation of the protein, namely C-terminal to N-terminal or the reverse. In the present case, additionally it is useful to circumvent edge effects due to filling of the pore at the front end and emptying at the back end.
5. Notice that almost all of the data processing can be done offline.
6. The procedure is non-destructive. By detaching conjugates after each cycle in the serial version or at the end in the parallel version the sample can be fully recovered. Thus the procedure has an archival character. (As is common in such studies it is assumed that in the serial as well as parallel case there are no losses due to adsorption, washing, and other related experimental procedures used.)
7. It is assumed that the presence of conjugates does not adversely affect the unidirectional and approximately constant translocation rate properties displayed in [22]. In particular conjugate molecules may be designed to be charge-neutral.
8. The proposed procedure can be used for protein identification by using only a small number (2 to 4) of AA types for detection, as is done, for example, in [21]. In general identification can be done in much less time than sequencing.
9. With suitable changes the procedure may be adapted for the detection of post-translational modifications (PTMs) in proteins. For example, a PTM behaves like a conjugate attached to an AA as it adds volume to the latter. An unconjugated sequence may contain clues to the position and identity of the PTM.

## Appendix Conjugated sequence matrix for 6 proteins from the human proteome

The table below gives id information for six proteins from the human proteome (Uniprot id UP000005640_9606).

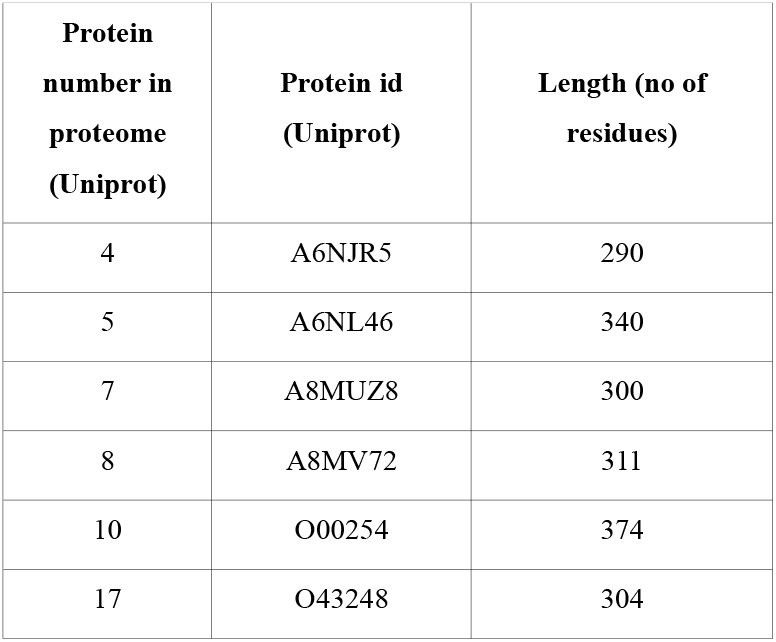

The following table shows the matrix of conjugated sequences containing the positions of occurrence of the 20 standard AAs in the first 100 residues of the above 6 proteins. (Only 100 residues are used to avoid data clutter and display problems.)

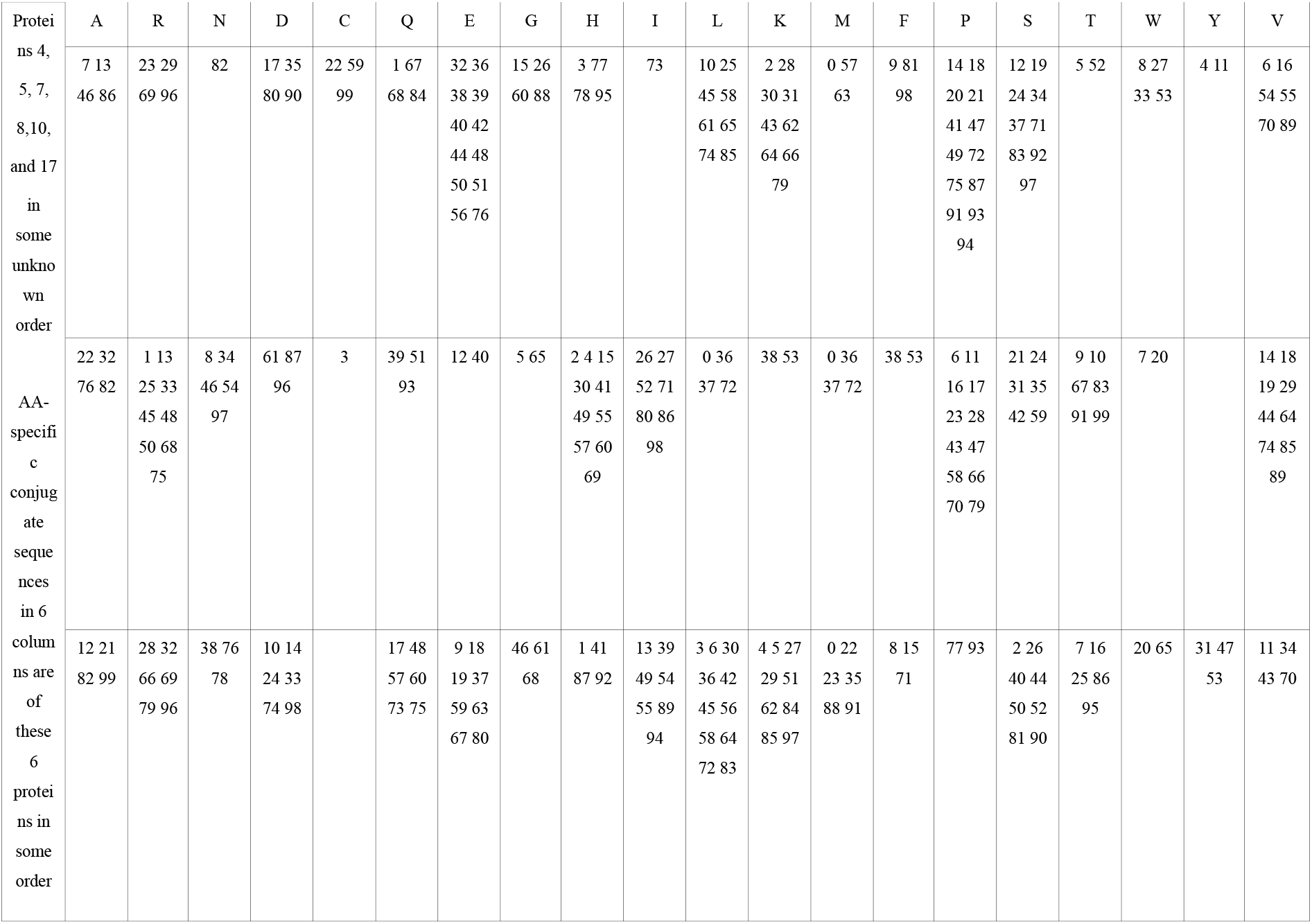
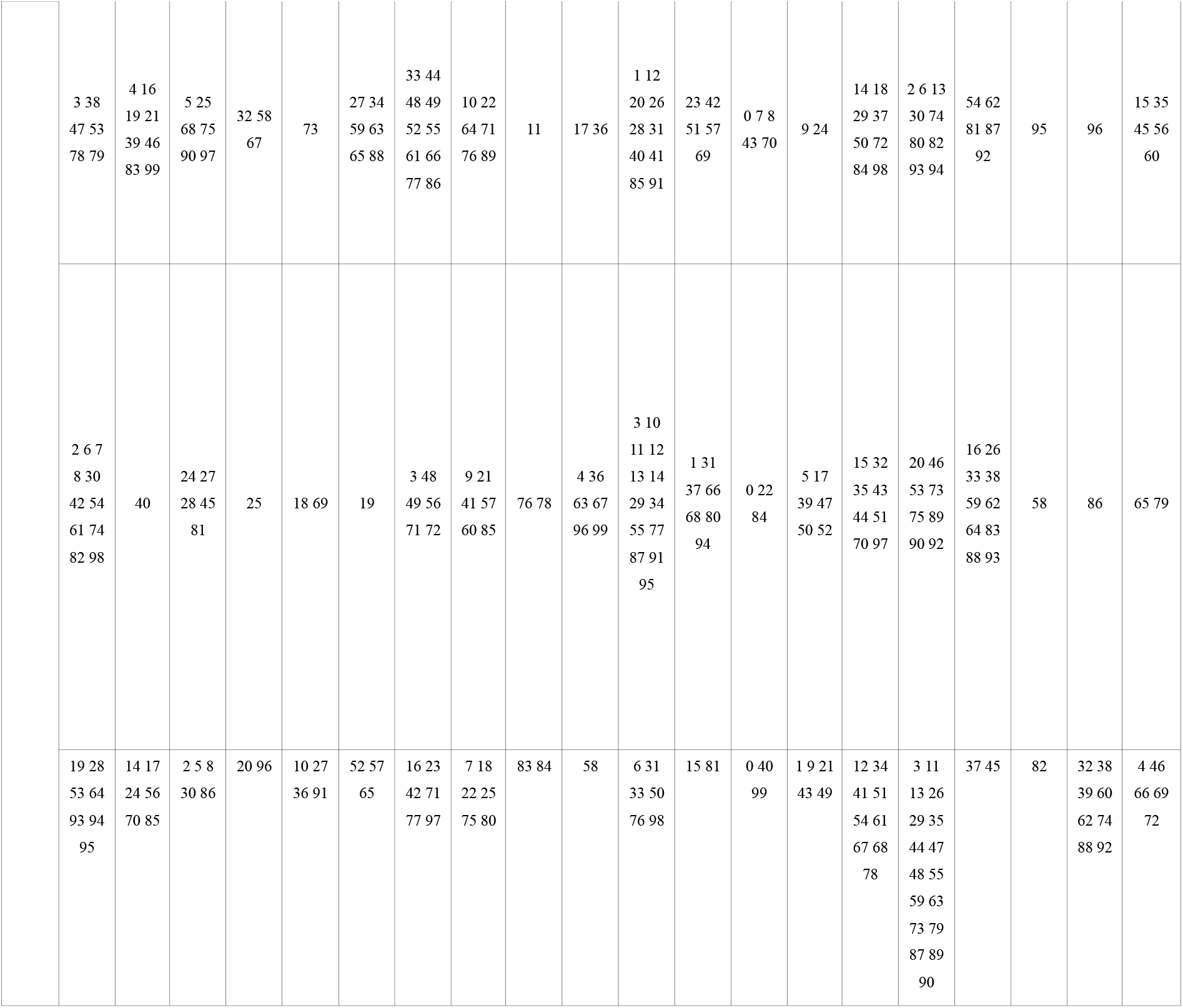

